# A method to quantitate maternal transcripts localized in sea urchin egg cortex by RT-qPCR with accurate normalization

**DOI:** 10.1101/2021.11.18.469148

**Authors:** Yulia O. Kipryushina, Mariia A. Maiorova, Konstantin V. Yakovlev

## Abstract

The sea urchin egg cortex is a peripheral region of eggs consisting of cell membrane and adjacent cytoplasm, which contains actin and tubulin cytoskeleton, cortical granules and some proteins required for early development. Method for isolation of cortices from sea urchin eggs and early embryos has been developed in 70s of 20th Century. Since that time this method has been reliable tool to study protein localization and cytoskeletal organization in cortex of unfertilized eggs and embryos during first cleavages. This study is an estimation of reliability of RT-qPCR to analyze levels of maternal transcripts that are localized in egg cortex. Firstly, we selected seven potential reference genes, 28S, *Cycb, Ebr1, GAPDH, Hmg1, Smtnl1* and *Ubb*, which transcripts are maternally deposited in sea urchin eggs. The candidate reference genes were ranked by five different algorithms (BestKeeper, CV, ΔCt, geNorm and NormFinder) upon calculated level stability in both eggs and isolated cortices. Our results show that gene ranking differs in total RNA and mRNA samples, though *Ubb* is most suitable reference gene in both cases. To validate feasibility of comparative analysis of eggs and isolated egg cortices by RT-qPCR, we selected *Daglb-2* as a gene of interest, which transcripts potentially localized in cortex, and found increased level of *Daglb-2* in egg cortices. This suggests that proposed RNA isolation method with subsequent quantitative RT-qPCR analysis can be used to approve cortical association of transcripts in sea urchin eggs.

## Introduction

RT-qPCR is a powerful tool to quantify gene expression levels in development, after exposure to chemical or physical treatment of cells *in vitro* and in many human deceases. Data normalization in RT-qPCR analysis is aimed to minimize errors in estimation of target mRNA levels. The most common approach is usage of endogenous reference genes (1). Perfect reference gene should have constant expression, while expression levels of many genes may be considerably changed during development and reveal different expression levels in different tissues and organs. So, each particular case requires seeking for reference genes that are most stably transcribed in all experimental samples. Using variably expressed genes as reference lead to obtaining incorrect results (2). Reference genes are chosen by comprehensive evaluation of gene expression stability of candidate genes by combinations of several methods.

Egg is a single cell, which has potential to develop into multicellular organism. Oocytes and eggs are polarized by asymmetrically deposited organelles and molecules within cytoplasm. Asymmetrically distributed maternal molecules, both RNAs and proteins, are key regulators of cell specification during early development. One of ancient mechanisms governing cell polarization is associated with localized RNAs found in oocytes of many model animals, like ascidians, *Drosophila*, zebrafish and *Xenopus*. Localized RNAs are also found in somatic cells, like neurons, oligodendrocytes, myoblasts, fibroblasts and epithelial cells(3, 4). In oocytes, different types of cytoskeleton play major role in anchoring of RNAs (5). Drosophila *nanos* is accumulated by diffusion and entrapment posteriorly by binding to actin filaments (6). *gurken* localization require static anchoring by Dynein at dorsal-anterior oocyte region and *oskar* posterior accumulation depends on its interaction with Kinesin heavy chain (7, 8). In Xeno*pus* oocytes, *Vg1* RNA is actively transported along microtubes and anchored to actin microfilaments in vegetal oocyte cortex (9).

In sea urchin eggs, cortex may play prevalence role for accumulation maternal factors that lead to establishment of polarity along both animal-vegetal and dorsal-ventral axes (10–12). Dishwelled, a protein of the Wnt/β-cathenin pathway regulating specification of vegetal blastomeres, which found in vegetal part of the eggs joined with egg cortex (13). Also, two mRNAs, *Panda* and *Coup-TF* found in subcortical area of oocytes, unfertilized eggs and early embryos. Subcortical *Panda* reveal gradient distribution which require to restrict Nodal signaling, which lead to dorsal-ventral axis formation in the sea urchin embryos (14). Coup-TF is a member of steroid-thyroid-retinoic acid superfamily, which control proper cell specialization along both animal-vegetal and dorsal-ventral axes. *Coup-TF* knockdown leads to lack nervous and digestive systems and ciliary band in embryos (15). Unequal distribution of maternal *Coup-TF* mRNA was detected not in all sea urchin species. *Coup-TF* were found to be localized laterally to animal-vegetal and 45° angle to dorsal-ventral axes in eggs of *Strongylocentrotus purpuratus* and *Lytechinus variegatus*, but not of *Paracentrotus redivivus* (16, 17). Some proteins necessary for development are associated with egg cortex reveal cortical distribution, which irrespective on directions of prospective developmental axes. Seawi and Vasa have been found in granules localized in egg cortex and later accumulated in primordial germ cells of sea urchin embryos (18, 19). Besides specified animal-vegetal and dorsal-ventral axes in sea urchin eggs early segregation of apical and basolateral cortical regions with involvement of Par proteins (20) suggest the presence in egg cortical region other unknown maternal factors that are necessary for epithelial organization of blastoderm.

Cell specification along embryonic axes and establishment of architecture of embryonic cells require many unequally distributed maternal factors in oocytes and eggs, many of them are still unknown for sea urchins. Exciting approach for quantitative RNA measurement called qPCR tomography was designed on *Xenopus* oocytes (21, 22). Principles of this method consist of RT-qPCR with RNA samples isolated from cryosections of oocytes that given along animal-vegetal axis. Thus, the authors propose to use qPCR tomography to analyze spatial expression patterns of RNAs in *Xenopus* oocytes localized in animal and vegetal poles. Availability of appropriate methods is good prerequisite for further development of methods to study spatial distribution of maternal transcripts in sea urchin oocytes and eggs. One of perspective approaches is comparative quantitative analysis of RNAs from sea urchin eggs and their isolated cortices. Method for cortex isolation has been employed for sea urchin eggs and embryos since 70s of 20th Century (23), and now we examine this method to isolate RNA with following RT-qPCR, which allows to measure levels of cortex-associated maternal transcripts.

The primary goals of this study are to evaluate suitability of quantitative RT-qPCR analysis of egg cortex-associated maternal transcripts and find appropriate reference genes for accurate signal normalization. Firstly, we found that RNA isolation from egg cortices is feasible procedure with additional stages to concentrate RNA samples. We selected some previously known (28S, *GAPDH, Hmg1* and *Ubb*), and found several new (*Cycb, Ebr1* and *Smtnl1*) candidate reference genes and performed RT-qPCR analysis of egg and isolated egg cortices. This set of genes was subjected to expression stability analysis by using BestKeeper (24), coefficient of variation (CV) (25), ΔCt (26), geNorm (27) and NormFinder (28) methods. We found that stability of most selected genes diverged in total RNA and poly(A) RNA samples. The highest stability in both cases was for *Ubb*, encoded polyubiquitin. So, this is the most suitable reference gene for comparative analysis of egg and cortex samples. Further, we predicted cortical localization of *Daglb-2* by transcriptome analysis and analyzed its levels in the eggs and cortices by RT-qPCR. We compared expression levels in both total and mRNA samples and found that increased level of *Daglb-2* was detected in mRNA samples of isolated cortices. This suggests that usage of mRNA fractions is successful to approve cortical association of *Daglb-2* in sea urchin eggs. Our results demonstrate a possibility to perform RT-qPCR analysis of isolated sea urchin egg cortices with accurate signal intensity normalization.

## Materials and Methods

### Animals and sample preparations

Adult *S. intermedius* sea urchins were collected in the Peter the Great Bay (Sea of Japan), kept in tubes with aerated sea water and fed with algae (*Ulva fenestrata* and *Saccharina japonica*) and carrot. Eggs were obtained by injection with 0.5M KCl. Eggs were washed several times with filtered sea water and then two times with CFSS (12mM HEPES, pH 7.6–7.8, 385mM NaCl, 10mM KCl, 21mM Na_2_SO_4_, 17mM glucose, 2.5mM MgCl_2_). Isolated cortices were prepared as described previously (19, 29, 30). Briefly, eggs attached to poly-L-lysine-coated coverslips (24×24 mm) were washed with CFSS supplemented with 5 mM EGTA. The coverslips were then gently washed by direct sprinkling with cortex isolation buffer (0.8 M mannitol) to remove majority of the egg content. Cortex samples were immediately used for RNA isolation. Isolated cortices were prepared on 6-8 coverslips for one RNA isolation.

### RNA extraction and cDNA synthesis

Total RNA was extracted from unfertilized eggs and isolated egg cortices using PureLink Mini kit (Thermo Fisher Scientific, USA) with some modifications. Total RNA from eggs was isolated according manufacturer’s manual from 4-5 μl of egg suspension. RNA from cortices were isolated using 2.5 ml of lysis buffer per 6-8 coverslips. Each coverslip was consequently placed in Petri dish filled with lysis buffer and cortices were lysed by pipetting. Then, the content was centrifuged through one spin cartridge after addition of equal volume of ethanol. To analyze total RNA, the samples were subsequently concentrated by GeneJet RNA Cleanup and Concentration Micro kit (Thermo Fisher Scientific, USA). Concentration was estimated by absorbance at 260 nm on Biophotometer (Eppendorf, Germany). Only samples with high purity (1.8-2.0 by ratio A260/A280) were used in analysis. To obtain poly(A) mRNA fraction, the RNA samples isolated by PureLink Mini kit were subsequently purified by Magnetic mRNA Isolation kit (New England Biolabs, USA). mRNA concentration was estimated by Qubit RNA HS Assay Kit (Thermo Fisher Scientific, USA). The first strand cDNA was synthetized using ProtoScript II kit (New England Biolabs, USA) from 1 μg of total RNA or 1.5 ng of mRNA with Random Primer Mix (2 and 0.5 μl, respectively).

### Selection of candidate reference genes and gene of interest

28S rRNA gene and three protein-coding genes, *GAPDH*, *Hmg1*, *Ubb*, were previously used as reference genes for embryonic and adult samples of different sea urchin species (31–34). To found new candidate genes, we analyzed *de novo* assembled transcriptome (35) using SRAs obtained from entire eggs and isolated cortices (BioProject PRJNA686841). Transcriptome assembly was performed by Trinity using Galaxy web-based platform (36) (https://usegalaxy.org). Three protein-coding genes, *Cycb, Ebr1* and *Smtnl1*, were selected from list of genes upon preliminary differential expression analysis (37). *Cycb, Ebr1* and *Smtnl1* transcripts are abundant and their FPKM values are not significantly differ in samples of eggs and isolated cortices (Table 1). Gene of interest, *Daglb-2*, was selected from list of cortically enriched transcripts, which FPKM values is >0.5 and significantly higher in cortices (Table 1). All used protein-coding sequences were found in the transcriptome. A part of 28S sequence were amplified and sequenced with primers designed to close species *S. purpuratus* (GenBank Ac. No. AF212171.1): Forward (CGCCCAACAGCTGACTCAGA) and Reverse (TAGCACCAGAAATCGGACGAA). All sequences were deposited in GenBank database. Accession numbers of sequences, primers and products’ sizes are done in Table 2.

**Table 1.** FPKM values of new candidate reference genes and gene of interest

| Gene | FPKM |  |
| --- | --- | --- |
|  | Eggs | Cortices |
| *Cycb* | 18086.99 | 15769.9 |
| *Daglb-2* | 2 | 9 |
| *Ebr1* | 7476.33 | 6490.31 |
| *Smtnl1* | 4183.87 | 3781.28 |

**Table 2.**
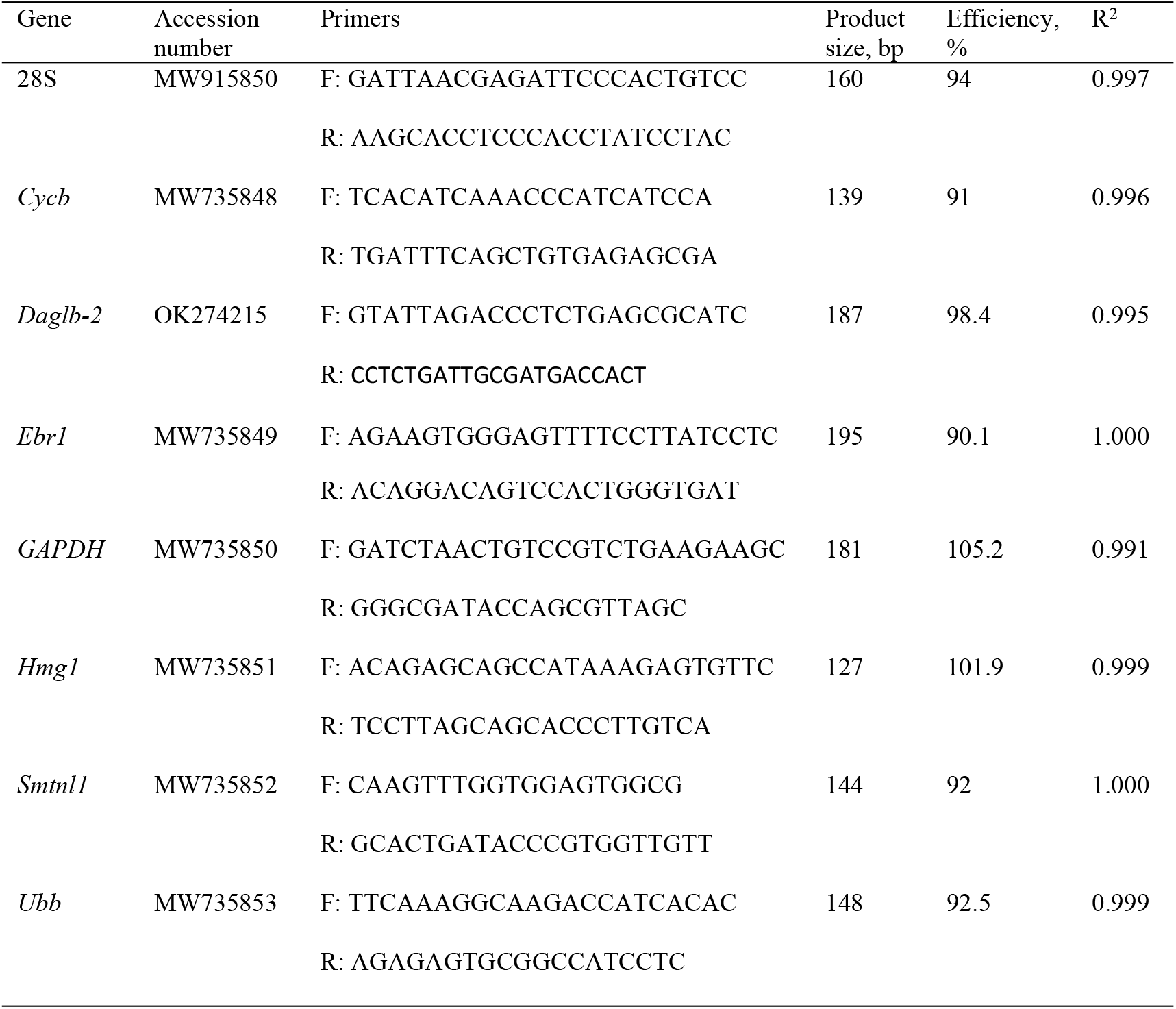
Names of genes, primers used for RT-qPCR and reaction efficiency.

### RT-qPCR

RT-qPCR was conducted on CFX96 Touch Real-Time PCR Detection System (Bio-Rad, USA) using qPCRmix-HS SYBR master mix (Evrogen, Russia) with the following temperature program: 94 °C for 30 sec, 40 cycles of 94 °C for 10 sec, 55 °C for 25 sec, 72 °C for 15 sec. After then, melting curve analysis was done. Three independent biological replicates were performed and each replicate was analyzed in technical triplicate. PCR efficiency was evaluated with the CFX Manager (Bio-Rad, USA).

### Data analysis

The stability of seven potential reference genes were analyzed using five approaches, BestKeeper (24), CV (25), ΔCt (26), geNorm (27) and NormFinder (28). *Daglb-2*, which potentially localized in egg cortex, was used to validate selected reference genes. Relative levels of *Daglb-2* in eggs and egg cortices were calculated according their Ct values using 2^-ΔΔCt^ method (38). Statistical analysis and data visualization were performed using GraphPad Prizm 9 Demo software (GraphPad Software, USA).

## Results

### Selection of candidate reference genes, specificity and amplification efficiency of RT-qPCR

Seven potential reference genes for comparative RT-qPCR analysis of eggs and isolated cortical layers were tested to find appropriate genes that can be used for accurate normalization. Four genes were selected based on literature data, 28S, *GAPDH*, *Hmg1*, *Ubb*. Three genes, *Cycb*, *Ebr1* and *Smtnl1*, were selected from list of preliminary tested genes that are abundant in both eggs and isolated cortices based on transcriptomics analysis (Table 1). Also, we tested several genes previously used for normalization or found by transcriptome analysis, but we omitted them because PCRs with our primers did not fit required amplification efficiency (90-110%).

All chosen genes were tested for reaction specificity which was determined by melting curve analysis. Single peaks were detected for all tested genes (Fig. S1). No signals were detected with all primer pairs without templates. Amplification efficiencies were calculated by standard curve method using two-, four- or five-fold serial dilutions of cDNA samples. Amplification efficiencies ranged between 90.1-105.2%. Correlation coefficients (R^2^) displayed values 0.991-1.000 (Table 2).

### Levels of candidate reference genes

Levels of tested candidate reference genes in six samples (three samples from eggs and three samples from isolated egg cortices) were evaluated by RT-qPCR in total RNA and mRNA samples. Raw and mean Ct values are given in Table S1. Maximum differences in Ct values between technical replicated were < 0.5 cycles. In total RNA samples, among tested genes 28S and *Ubb* were most abundant genes with lowest means of Ct values (15.6 and 15.91, respectively). *GAPDH* showed the lowest level with highest mean of Ct value (28.02). The highest Ct variation was for 28S (4.87 cycles) and the lowest one for *Smtnl1* (0.66 cycles) (Fig 1A). In mRNA samples, most abundant genes were *Ubb* (mean of Ct value 20.99) and 28S (mean of Ct value 21.52). *GAPDH* revealed minimal level with mean of Ct 32.24. 28S was most variative with Ct range of 3.79 cycles. The less variative gene was *Ubb* with Ct range of 1.1 cycles (Fig. 1B).

**Figure 1.**
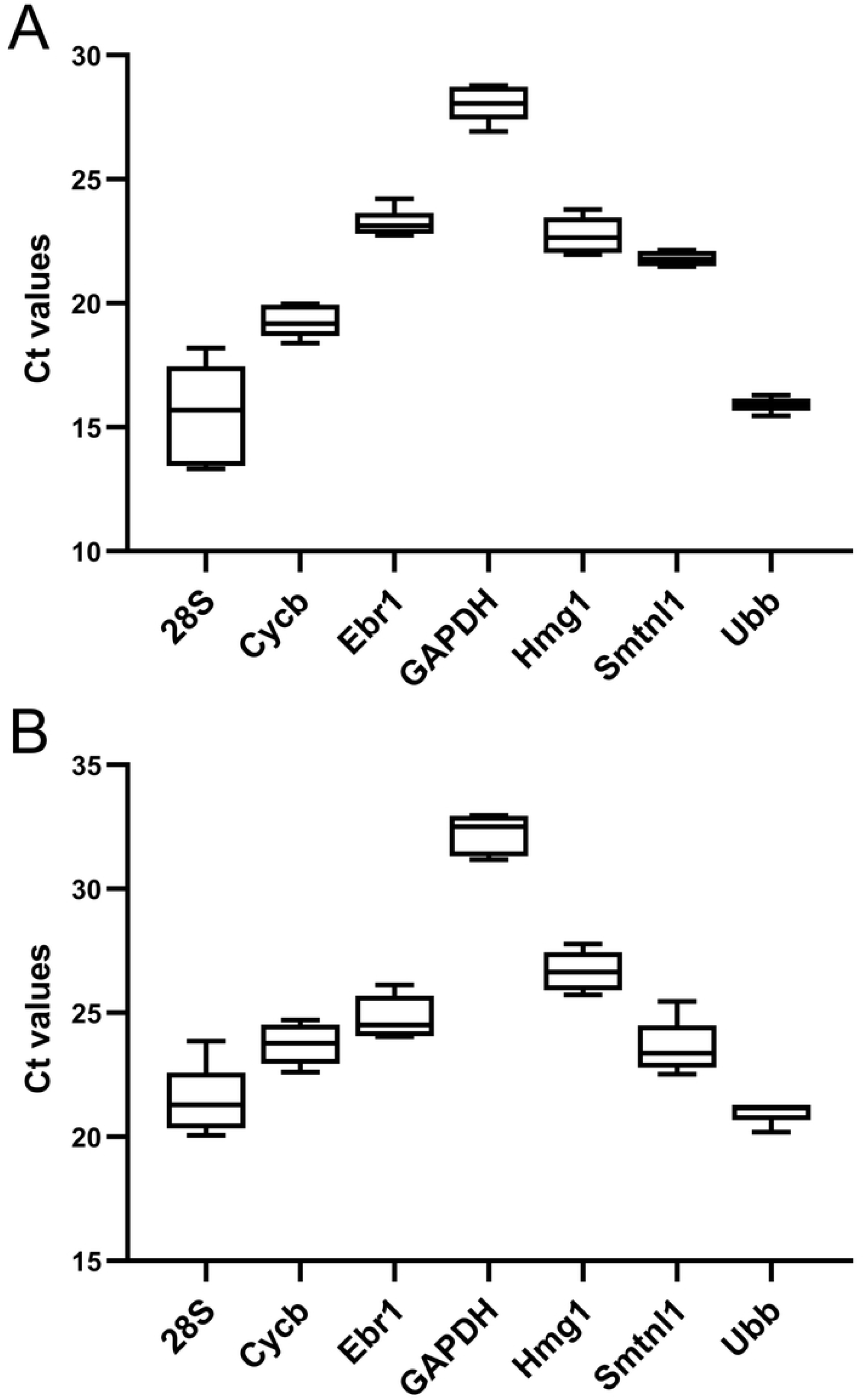
Boxplot of Ct values for candidate reference genes in all samples. (A) Ct values in cDNA samples synthetized from total RNA (B) Ct values in cDNA samples synthetized from mRNA. The boxes show interquartile range (25-75%), horizontal lines represent medians. The whiskers show the minimum and maximum values. No outliers were detected.

### Expression stability analysis and determination of minimal number of reference genes for normalization

We employed expression stability analysis using five different algorithms to evaluate level stability of transcripts in unfertilized eggs and egg cortices.

#### BestKeeper analysis

This method allows to analyze stability by SD and CV generated from raw Ct values (24). The lowest SD and CV values correspond to the highest stability. SD values lower than 1 indicate acceptance as reference genes. According to the SD values, the most stable gene for total RNA and mRNA samples was *Ubb* (SD value=0.21 and 0.32, respectively) (Fig. 2A). The least stable gene was 28S (SD=1.72 for total RNA and 0.99 for mRNA). Total RNA value is higher than 1, which is unacceptable for usage 28S as reference gene.

**Figure 2.**
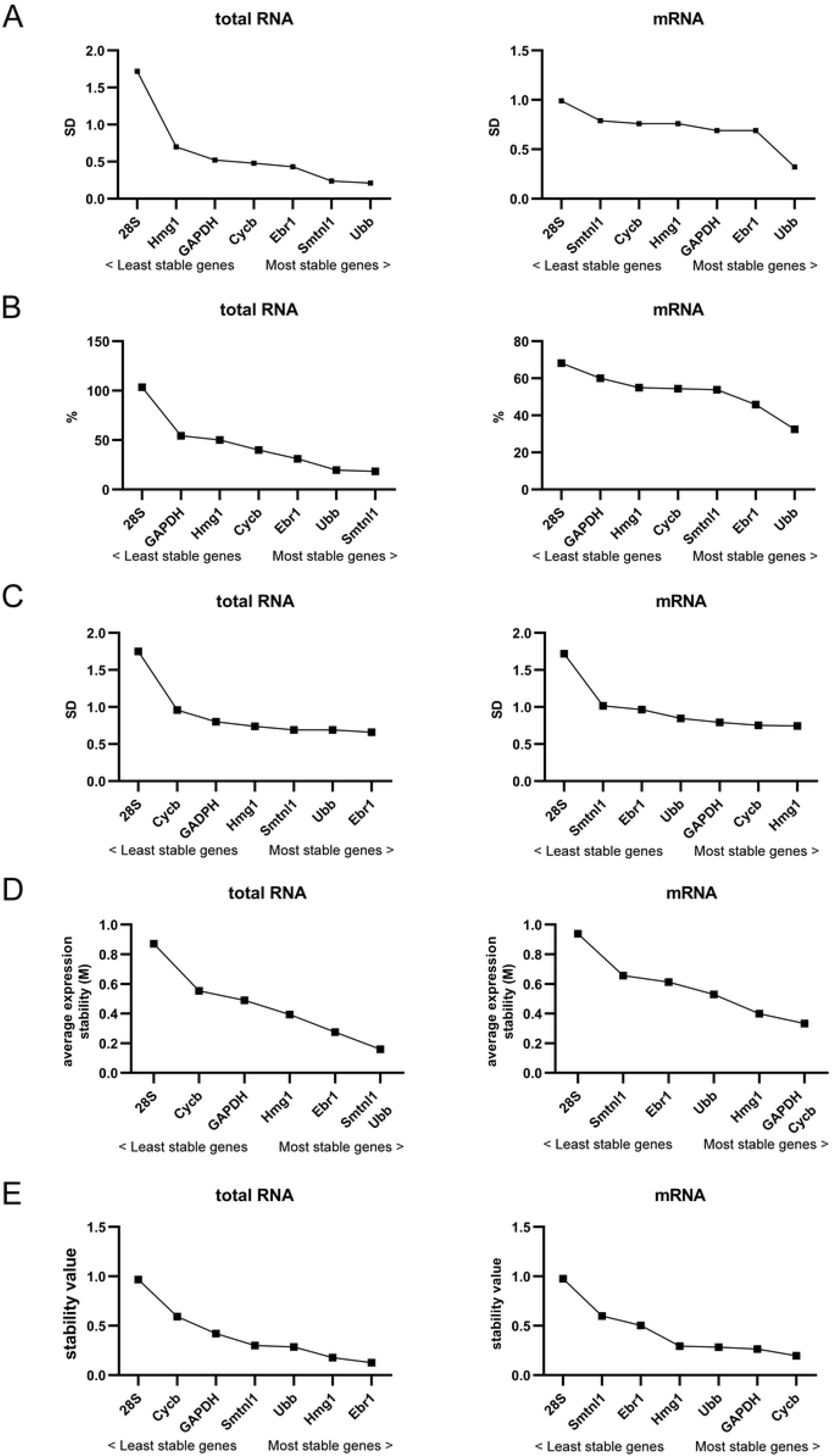
Stability analysis of candidate reference genes performed by different methods. Stability estimated by BestKeeper (A), CV (B) ΔCt (C), geNorm (D) and NormFinder (E).

#### CV analysis

CV method is simply based on comparison of CV of expression levels. The lowest CV value, which is defined as a ratio of SD to average 2^C_tmin_-C_tsample_^, corresponds to highest intragroup stability (25). The least stable gene in both total RNA and mRNA samples was 28S with CV values 103.4% and 68.17%, respectively. The most stable gene in total RNA samples was *Smtnl1* (18.41%). In mRNA samples, *Ubb* showed highest stability (32.43%) (Fig. 2B).

#### ΔCt analysis

The Δ Ct method is based on pairwise comparisons and calculation SD of ΔCt values for each pair of genes (26). The lowest value of average SD corresponds to the highest stability of expression among evaluated genes. Our results showed that for total RNA the most stable gene was *Ebr1* (0.66) and the least stable 28S (1.75) (Fig. 2C). In mRNA samples, the most stable gene was *Hmg1* (0.743) and least stable was also 28S (1.72).

#### geNorm analysis

This method is based on pairwise variation that consequently exclude least stable genes after each step of analysis. Finally, two most stable genes are determined. geNorm utilize average expression stability (M) values (27). Threshold M value is below 0.5 indicate good reference genes. The most stable genes have the lowest M values. As shown, among seven tested genes the best pair for total RNA was *Smtnl1/Ubb* with M value 0.16 (Fig. 2D). 28S and *Cycb* showed M values above 0.5 that indicate their inapplicability as reference genes. The best pair in mRNA analysis was *GAPDH/Cycb* with M value 0.33. on the second place was *Hmg1* (M value 0.4). Values of other genes were above 0.5 which inappropriate for good reference genes. Among them, 28S was the least stable (M value 0.94) (Fig. 2D). Another parameter calculated by geNorm is pairwise variation (V_n/n+1_) between normalization factors. It allows to define minimal number of reference genes for accurate normalization. Cut-off threshold 0.15 is recommended to find out the optimal number reference genes (27, 39). In our test, all pairwise variations in both total RNA and mRNA cases were below 0.15 (Fig. 3) that point to usage of two reference genes for normalization.

**Figure 3.**
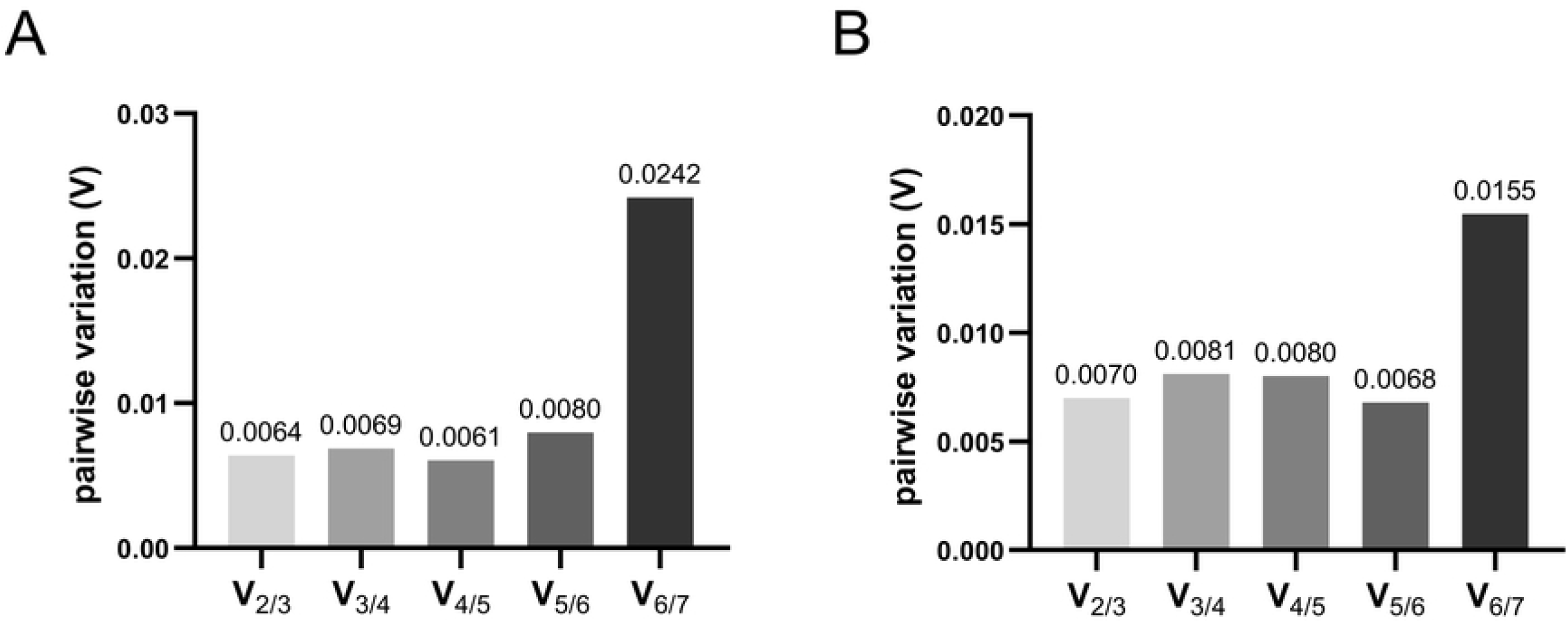
Determination of minimal number of reference genes for accurate normalization. Pairwise variations (Vn/n+1) were calculated by geNorm in all samples (eggs and isolated egg cortices) for total RNA (A) and mRNA (B).

#### NormFinder analysis

This method takes into account both intragroup and intergroup expression variability. The most stable genes have lowest stability values (28). NormFinder analysis showed that the most stable gene was *Ebr1* (0.125) and the least 28S (0.97) in total RNA analysis (Fig. 2E). Additionally, NormFinder determined the *Ebr1*/*Hmg1* pair as the best combination of reference genes, as these genes have lowest values. In mRNA samples, *Cycb* revealed highest stability (0.2) and 28S the lowest one (0.98). The best pair of reference genes NormFinder proposed *Cycb*/*GAPDH*.

After analysis by different methods, we summarized ranking of candidate reference genes, which is presented in Table 3. For total RNA, general view showed that *Ubb*, *Ebr1* and *Smtnl1* are three most stable, therefore they are most appropriate reference genes. BestKeeper analysis revealed that *Ubb* was most stable gene. ΔCt and NormFinder analysis detected *Ebr1* as most stable and according to CV method the most stable gene was *Smtnl1*. geNorm does not allow to recognize the best reference gene, this method determine the pair of genes with highest stability. For total RNA the best pair of reference genes was found to be *Smtnl1/Ubb*. Calculations of data obtained from mRNA samples showed other ranking of genes (Table 3). *Ubb* was ranked as most stabe by BestKeeper and CV methods. ΔCt determined *Hmg1* as most stable gene, and *Cycb* was most appropriate reference gene in NormFinder analysis. geNorm found *Cycb*/*GAPDH* as most stable pair. The only one gene, *Ubb*, was found among the most stable genes in two our separate analyses of total RNA and mRNA. 28S ranked as least stable gene in analyses on both type of RNA samples.

**Table 3.**
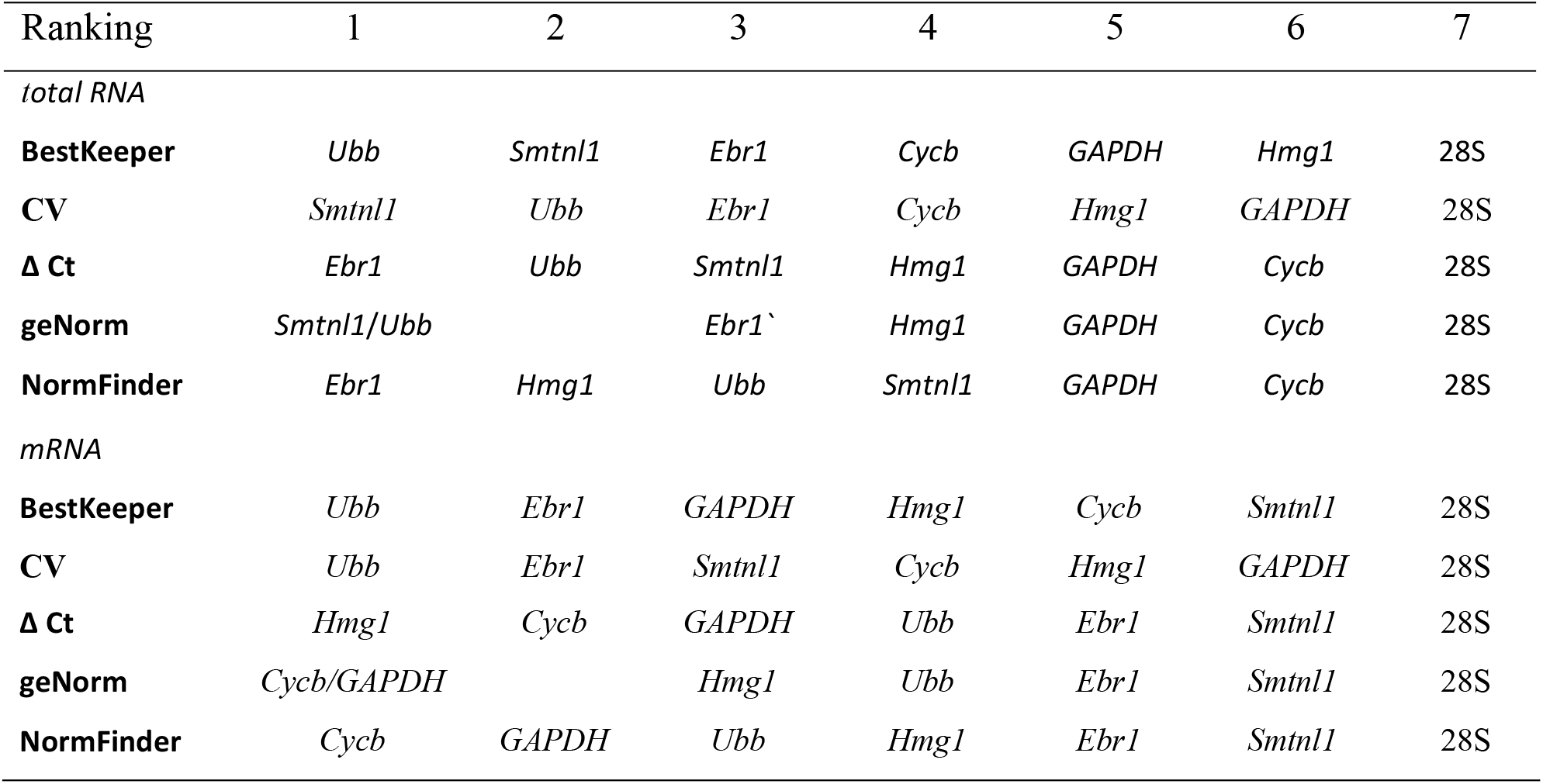
Stability ranking of candidate reference genes given upon BestKeeper, CV, Δ Ct, geNorm and NormFinder.

| Ranking | 1 | 2 | 3 | 4 | 5 | 6 | 7 |
| --- | --- | --- | --- | --- | --- | --- | --- |
| *total RNA* |  |  |  |  |  |  |  |
| **BestKeeper** | *Ubb* | *Smtnl1* | *Ebr1* | *Cycb* | *GAPDH* | *Hmg1* | 28S |
| **CV** | *Smtnl1* | *Ubb* | *Ebr1* | *Cycb* | *Hmg1* | *GAPDH* | 28S |
| **$\Delta$ Ct** | *Ebr1* | *Ubb* | *Smtnl1* | *Hmg1* | *GAPDH* | *Cycb* | 28S |
| **geNorm** | *Smtnl1/Ubb* |  | *Ebr1`* | *Hmg1* | *GAPDH* | *Cycb* | 28S |
| **NormFinder** | *Ebr1* | *Hmg1* | *Ubb* | *Smtnl1* | *GAPDH* | *Cycb* | 28S |
| *mRNA* |  |  |  |  |  |  |  |
| **BestKeeper** | *Ubb* | *Ebr1* | *GAPDH* | *Hmg1* | *Cycb* | *Smtnl1* | 28S |
| **CV** | *Ubb* | *Ebr1* | *Smtnl1* | *Cycb* | *Hmg1* | *GAPDH* | 28S |
| **$\Delta$ Ct** | *Hmg1* | *Cycb* | *GAPDH* | *Ubb* | *Ebr1* | *Smtnl1* | 28S |
| **geNorm** | *Cycb/GAPDH* |  | *Hmg1* | *Ubb* | *Ebr1* | *Smtnl1* | 28S |
| **NormFinder** | *Cycb* | *GAPDH* | *Ubb* | *Hmg1* | *Ebr1* | *Smtnl1* | 28S |

### Validation of candidate reference genes

To validate the reliability of recommended reference genes, *Daglb-2* was selected from list of cortically-enriched transcripts. Normalization of signal intensity was done using top-ranked genes and least stable 28S. *Ebr1, Smtnl1* and *Ubb* were found to be most appropriate reference genes for total RNA samples (Table 3). After normalization to *Ebr1, Smtnl1, Ubb* and pair *Smtnl1/Ubb* reference genes, the levels of *Daglb-2* in egg cortices were little bit lower than in eggs (Fig. 4A), but these differences between samples and reference genes were statistically insignificant. Normalization to the least stable 28S gene showed 5.64-fold higher level of *Daglb-2* in cortices. By using total RNA samples, we did not support that *Daglb-2* is cortically-enriched transcript. Nevertheless, nearly equal *Daglb-2* signals normalized to the top-ranked genes in cortices indicate the reliability of chosen reference genes. *Daglb-2* levels measured in mRNA samples were normalized to appropriate top-ranked genes, *Cycb*, *Hmg1*, *Ubb* and *Cycb*/*GAPDH* (Table 3). In contrast to values calculated in total RNA samples, mRNA analysis showed increased levels of *Daglb-2* in cortices (Fig. 4A), from 2.18-fold higher level in case of *Cycb* to 2.65-fold higher level in case of *Ubb*. Although values normalized to *Cycb*, *Hmg1* and *Cycb*/*GAPDH* in cortices were higher than in eggs, they did not reveal statistical significance, which may ensure from variable Ct values of these genes (Fig. 1B). Only values normalized to *Ubb*, which Ct were in narrow range, were statistically significant (Fig. 4A). We compared relative levels of *Daglb-2* normalized to *Ubb* in total RNA and mRNA (Fig. 4B). Total RNA samples did not detect significant difference in mRNA levels between eggs and isolated cortices, while purified mRNA allowed to detect significant enrichment of *Daglb-2* in cortices. It suggests that analysis of mRNA is more sensitive than total RNA. 28S which is a worst reference gene in both types of samples showed high increasing of *Daglb-2* levels in cortices due to low levels in cortices.

**Figure 4.**
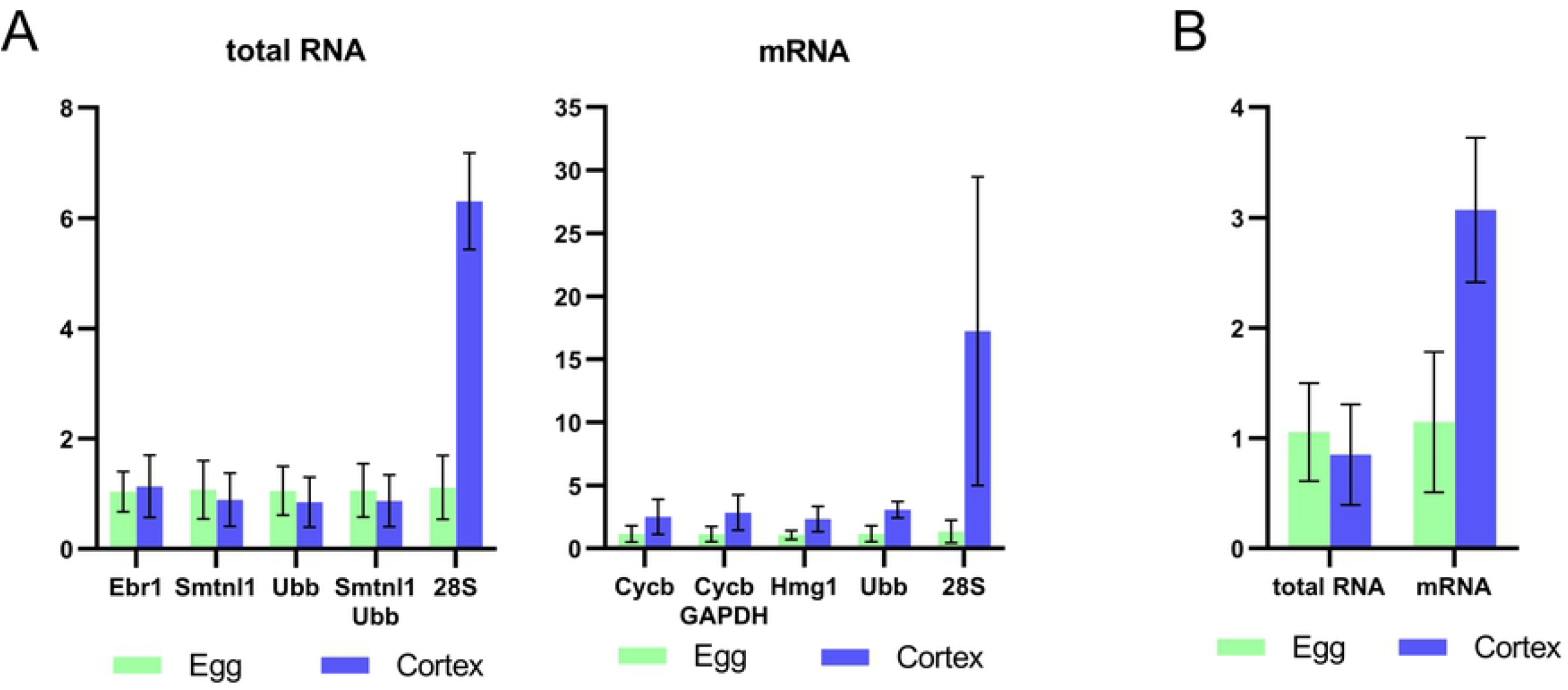
Comparison of the normalized relative levels of *Daglb-2* between eggs and isolated cortices. (A) To normalize values, three most stable genes and best pair of genes were selected. Also, data were normalized to least stable gene (28S). (B) Relative levels of *Daglb-2* normalized to *Ubb*.

## Discussion

RT-qPCR is convenient method to estimate gene expression in different biological contexts. The primarily goal of this study is to design approach based on RT-qPCR for quantitative analysis of cortically-associated maternal transcripts of sea urchin eggs. The proposed approach is based on comparison the levels of genes of interest between eggs and isolated egg cortices. The first necessary prerequisite to perform this analysis is a suitable method for RNA isolation. We adapted column-based RNA isolation protocol for isolated egg cortices. Isolated total RNA may be subsequently processed to obtain purified poly(A) RNA. Second prerequisite for accurate RT-qPCR analysis is usage of appropriate reference transcripts (genes), which levels are less variable among eggs and isolated cortices. We analyzed four candidate reference genes selected from known reference genes and three relatively abundant transcripts detected in both entire eggs of *S. intermedius* and their isolated cortices. To evaluate comprehensively the stability of selected genes, 28S, *GAPDH*, *Hmg1*, *Ubb*, *Cycb*, *Ebr1* and *Smtnl1*, we used five different methods and compared derived results. During analysis of total RNA samples, three programs, BestKeper, Δ Ct and geNorm, showed similar results. According to them, *Ebr1*, *Smtnl1* and *Ubb* are three most stable genes (Table 3). Each program ranked these genes differently, but in each case these genes were included in top 3 as most stable. Results of NormFinder were different, *Ebr1* was most stable gene, *Hmg1* and *Ubb* were ranked as second and third stable genes, respectively. We decided to designate *Ebr1*, *Ubb* and *Smtnl1* as most suitable reference genes upon analysis by BestKeper, Δ Ct and geNorm. Estimation of level stability in mRNA samples ranged candidate reference genes differently than in total RNA samples. According to BestKeeper and CV methods *Ubb* is most stable gene, while other methods showed *Hmg1* (Δ Ct), *Cycb* (NormFinder) and *Cycb/GAPDH* pair (geNorm) as best reference genes.

Among all tested genes, new and previously reported as internal controls (31–34, 40, 41), the only *Ubb* showed suitability for quantitative RNA analysis of total RNA and mRNA samples from isolated egg cortices. *Ubb* encodes polyubiquitin is well-known reference gene in quantitative expression analysis of sea urchin embryos, as level of *Ubb* mRNA is relatively stable in sea urchin development (31, 42). Also, *Ubb* is suitable for both qualitative and quantitative expression analysis during sea urchin gametogenesis (43, 44). Two new candidate reference genes found by our transcriptomic analysis as abundant transcripts in both eggs and egg cortices, *Ebr1* and *Smtnl1*, previously have not been used as internal control. These two genes were shown to be suitable only for total RNA. *Ebr1* encodes egg cell-surface protein. It is one of proteins that responsible for species-specific sperm adhesion to sea urchin eggs via interaction with Bindin localized on spermatozoan surface (45, 46). *Smtnl1* encodes a muscle protein participating in regulation of muscle contraction and adaptation in mammals (47), while in sea urchin embryos its functions remain unstudied.

A main goal of this study is testing reliability of quantitative approach to analyze transcripts, which anchored in subcortical area of eggs. To test reliability our approach, we should use as gene of interest transcript potentially associated with egg cortex. We excluded from our analysis transcripts have found to be localized presumably in cortex, *Panda* and *Coup-TF* (14, 17). We have not found *Panda* homolog in *S. intermedius* egg transcriptome. Assembled part of *Coup-TF* is GC-rich region, which poorly amplified by RT-qPCR. To find cortex-associated transcript for our analysis, we selected transcripts, which levels were higher in cortices that in eggs. *Daglb-2* was chosen as transcript, which may be localized in egg cortex according to transcriptomic analysis. *Daglb-2* encodes homolog of mammal transmembrane enzyme diacylglycerol lipase beta localized in plasma membrane, where it generates endocannabinoids from membrane lipids. Endocannabinoids participate in inflammatory response in macrophages and act as neurotransmitters in central and peripheral nervous systems (48–50). Probably, *Daglb-2* localization in egg cortex requires for local translation of the encoded protein, which is integrated in plasma membrane. However, biological role of *Daglb-2* for sea urchin embryogenesis is still unclear. Upon our transcriptomic analysis and predicted functions of *Daglb-2* we think that this gene is suitable as gene of interest to approbate approach of quantitative evaluation of cortex-associated transcripts in sea urchin eggs.

Many researchers normalize signal intensities of studies genes against single reference gene. Nevertheless, it is necessary to confirm invariant expression of potential reference gene under all experimental conditions. Alternatively, usage of two or more reference genes is the better choice (27, 51). geNorm allows to determine number of reference genes for reliable normalization (two genes is minimal number). Our analysis showed Hnot significantly differ among either total RNA or mRNA samples. Fundamental differences were between total RNA and mRNA data. Total RNA samples did not show significant differences of *Daglb-2* levels between eggs and their cortices. Analysis of purified mRNA revealed increased *Daglb-2* in cortices supporting our differential expression results. While normalization to any best reference gene showed similar ratios of *Daglb-2* levels between eggs and cortices, only *Ubb* showed significant differences. Taking into account these results and lowest Ct variation of *Ubb* among evaluated genes, we propose that *Ubb* is the best choice for intensity signal normalization. *Ubb* was defined as the most stable gene by Best Keeper and CV methods and ΔCt, geNorm and NormFinder other methods ranked other genes on the first place. Different methods for evaluation of expression stability are based on different principles. They may well define the least stable gene, but the best genes are different (52). In case of total RNA samples, we propose to use any best gene, *Ubb*, *Ebr1* and *Smtnl*. Nevertheless, we found that RT-qPCR with total RNA samples had not been sensitive to detect differences of *Daglb-2* levels between eggs and cortices.

## Conclusions

This study provides the first report of RT-qPCR analysis of maternal transcript to verify its association with egg cortex in sea urchin eggs, which has been predicted by transcriptomic analysis. Firstly, optimization of RNA isolation method from isolated cortices show possibility to extract successful amount of RNA for cDNA synthesis. Next evaluation of possible reference genes allowed to find appropriate genes that can be used for signal intensity normalization either for total RNA and mRNA samples. Finally, RT-qPCR analysis of presumably cortex-associated *Daglb-2* showed that increased levels of its transcript in egg cortices has been only detected in mRNA samples. Thereby RT-qPCR may be utilized as one of methods of analysis of mRNA spatial distribution in sea urchin eggs.

## Acknowledgements

This work was supported by the Russian Foundation for Basic Research (Grant number: 20-04-00332). The authors are grateful to Dr. Andrey Kukhlevsky for Sanger sequencing.

## Supporting information captions

**Figure S1.** Melt curve analysis of PCR products from one biological replicate. Egg and cortex melt curves are given together for each gene. Negative control (template-free) is marked by Neg.

